# From pelagic to reef: Genomic basis of morphological adaption during life-history transition in the orbicular batfish (*Platax orbicularis*)

**DOI:** 10.64898/2026.09.10.750547

**Authors:** Yuxuan Zhang, Yanwen Shao, Liang Zhong, Runsheng Li, Wenlong Cai

**Author notes:** Corresponding authors: Runsheng Li and Wenlong Cai.

## Abstract

Metamorphosis enables teleosts to transition between ecological niches, driving lineage-specific morphological innovations. The orbicular batfish (*Platax orbicularis)* exemplifies this, transforming from a pelagic larva into a reef-associated juvenile with extreme allometric fin elongation. Here, we generated a chromosome-level genome assembly of 715.46 Mb, with 99.4% completeness. Integrating comparative genomics with tissue-specific transcriptomics, we demonstrate that such transformation is a systemic process shaped by both long-term evolutionary adaptation and short-term developmental reprogramming. Comparative analyses revealed expansion of development-related gene families and significant evolution of immune- and sensory-related families, likely associated with ecological niche transition. Spatiotemporal clustering of metamorphosing fin transcriptomes identified rapid cell cycle activation accompanied by sustained suppression of basal metabolism, indicating developmental trade-offs during fin elongation. Several differentially expressed genes involved in cell cycle (*ankfn1* and *mcph1*), ossification (*cyp24a1* and *sema4d*), and extracellular matrix remodeling (*sdc4* and *serpine1*) exhibited positive selection, suggesting coordinated regulatory and protein-level adaptation. Among an extensive repertoire of 49 *hox* genes, four (*hoxb6b*, *hoxc8a*, *hoxd9b*, *hoxd10a*) were differentially expressed, implicating developmental regulators in fin remodeling. Together, our findings offer new insights into the evolutionary context and molecular mechanisms of metamorphosis in orbicular batfish, and advance the understanding of adaptive evolution and metamorphosis innovations in teleosts.

## 1. Introduction

Marine teleosts generally possess a bipartite life history, involving a critical transition from pelagic larvae to demersal, benthic or still pelagic juveniles. This process is typically accompanied by morphological, physiological, and behavioral changes, known as metamorphosis ^1^. For example, as clownfish transition from pelagic larvae to reef-associated juveniles, they shift from an elongated to a more robust body shape, alongside pigmentation alterations and metabolic reprogramming^2,3^. Such remodeling is understood as an integrated developmental process that supports ecological adaption, which is crucial for individual survival and population persistence^4^.

Different teleost lineages have evolved lineage-specific metamorphic trajectories in response to ecological transitions^5^. In many species, these trajectories involve remarkable morphological innovations. For example, the elongated fins of larval groupers regress during their transition into reef-associated juveniles^6^, whereas soles experience eye migration together with asymmetrical cranial ossification and pigmentation as they settle into benthic habitats^7^. Conversely, taxa like seabreams undergo metamorphosis without profound morphological transformations^8^. These differences indicate that marine teleost metamorphosis is evolutionarily diverse, reflecting lineage-specific solutions to ecological challenges. However, the genomic basis underlying such lineage-specific innovations remains poorly understood.

Deciphering the molecular mechanisms of metamorphosis is essential for advancing the knowledge of these transformations in marine teleost development. For economically important aquaculture species, elucidating the molecular changes that accompany critical metamorphic stages may also help improve early survival and juvenile performance^9,10^. Previous studies have primarily focused on endocrine hormones and their related signaling pathways, establishing their core regulatory roles in developmental transitions and morphological remodeling^5,11^. For example, thyroid hormones (TH) are crucial for the transition from larvae to juveniles in clownfish, regulating both morphogenesis and metabolic reprogramming^3^. Similarly, in groupers, activation of TH and corticosteroid pathways have been associated with regression of transient larval spines^12^. Nevertheless, metamorphosis is not only an endocrine-driven process of body-wide transformation, but also involves organ- or tissue-specific remodeling and physiological reorganization that likely contribute to adaptation to the juvenile environment. To date, few studies have elucidated these organ- or tissue-specific downstream gene regulatory networks.

*Platax* spp., reef-associated fish belonging to the family Ephippidae and the order Acanthuriformes, provide an excellent system for investigating these questions. They are considered important contributors to the ecological balance of coral reef environments by preventing algal overgrowth despite their omnivorous nature^13^. The *Platax* species exhibit a remarkable metamorphosis accompanied by highly distinctive morphological transformations, for which this genus is particularly well known. Beginning life as pelagic larvae, these fish subsequently migrate towards coral reefs and mangrove areas, where they develop into juveniles^14^. During this transition, their body forms a deep and compressed shape^15^. Most notably, the dorsal, pelvic, and anal fins extremely elongate, reaching or even exceeding body depth, producing the characteristic bat-like appearance of juveniles^15,16^. The fin morphology is thought to be associated with swimming performance adapted to the hydrodynamic conditions of coral reef environments^17^. However, at the adult stage, the body of the orbicular batfish becomes more rounded, making its previously elongated fins appear shorter relative to the body. Currently, the *Platax* species has also become an economically valuable food fish because of their minimal bones and abundant meat^18^. However, despite their ecological and economic importance, research on *Platax* species has focus mainly on ecology and behavior, while the genomic basis and molecular programs associated with their unique metamorphic traits remain unknown.

In this study, we investigated the metamorphosis of orbicular batfish (*Platax orbicularis*), which is one of the most prominent species within the *Platax* genus. We generated the first chromosome-level genome for this species and investigated its lineage-specific adaptation through comparative analysis with other representative fish species. Furthermore, since dorsal, pelvic, and anal fins are the most prominently remodeled organs during the metamorphosis of orbicular batfish, we performed transcriptomic sequencing of these fins across the critical transition from larvae to juveniles, aiming to characterize the organ-specific gene regulatory networks driving this morphological remodeling. Overall, we provide the first genomic resource for orbicular batfish and offer insights into the molecular basis of ecological adaptation and tissue remodeling during metamorphosis in *Platax* fishes from both evolutionary and developmental perspectives. Importantly, our study facilitates the understanding of morphological plasticity and ecological adaptation during marine teleost development, and may also contribute to the improvement of aquaculture practices for *Platax* species as economic fish during their early developmental stages.

## 2. Methods

### 2.1 Ethics statement

All animal procedures were approved by the City University of Hong Kong Animal Care and Use Committee (AN-STA-00000890). We have complied with all relevant ethical regulations for animal use.

### 2.2 Sample collection

All orbicular batfish used in this study were obtained from the Hong Kong Agriculture, Fisheries and Conservation Department. One three-year-old individual (sex-undetermined) was used to generate DNA sequencing data for genome assembly. Muscle tissue from this fish was flash-frozen in liquid nitrogen and stored at −80LJ°C until DNA extraction. Another two-month-old individual was used to generate RNA sequencing data for gene annotation. Tissues from brain, gill, heart, head kidney, spleen, and intestine of this fish were collected and stored at −80LJ°C until RNA sequencing.

To investigate the gene regulatory networks involved in metamorphosis from larvae to juveniles, we performed RNA sequencing on orbicular batfish at different metamorphic stages. All individuals were approximately 25 days old, but were at different metamorphic stages because of variation in growth rate. Based on the morphological characteristics described by Leu, et al. ^16^, individuals were classified and collected at the pre-, mid-, and post-metamorphic stages from larva to juvenile (Fig. 1a). Before tissue collection, fish were euthanized with an overdose of MS-222 mixed with an equal amount of sodium bicarbonate. For each fish, we sampled and pooled the dorsal, pelvic, and anal fins for RNA-Seq, which undergo the most prominent changes during metamorphosis. At the pre-metamorphic stage, three biological replicates were prepared, each comprising pooled fin tissues from three fish. At the mid- and post-metamorphic stages, three biological replicates were prepared for each stage, with each replicate derived from a single fish.

**Figure 1.**
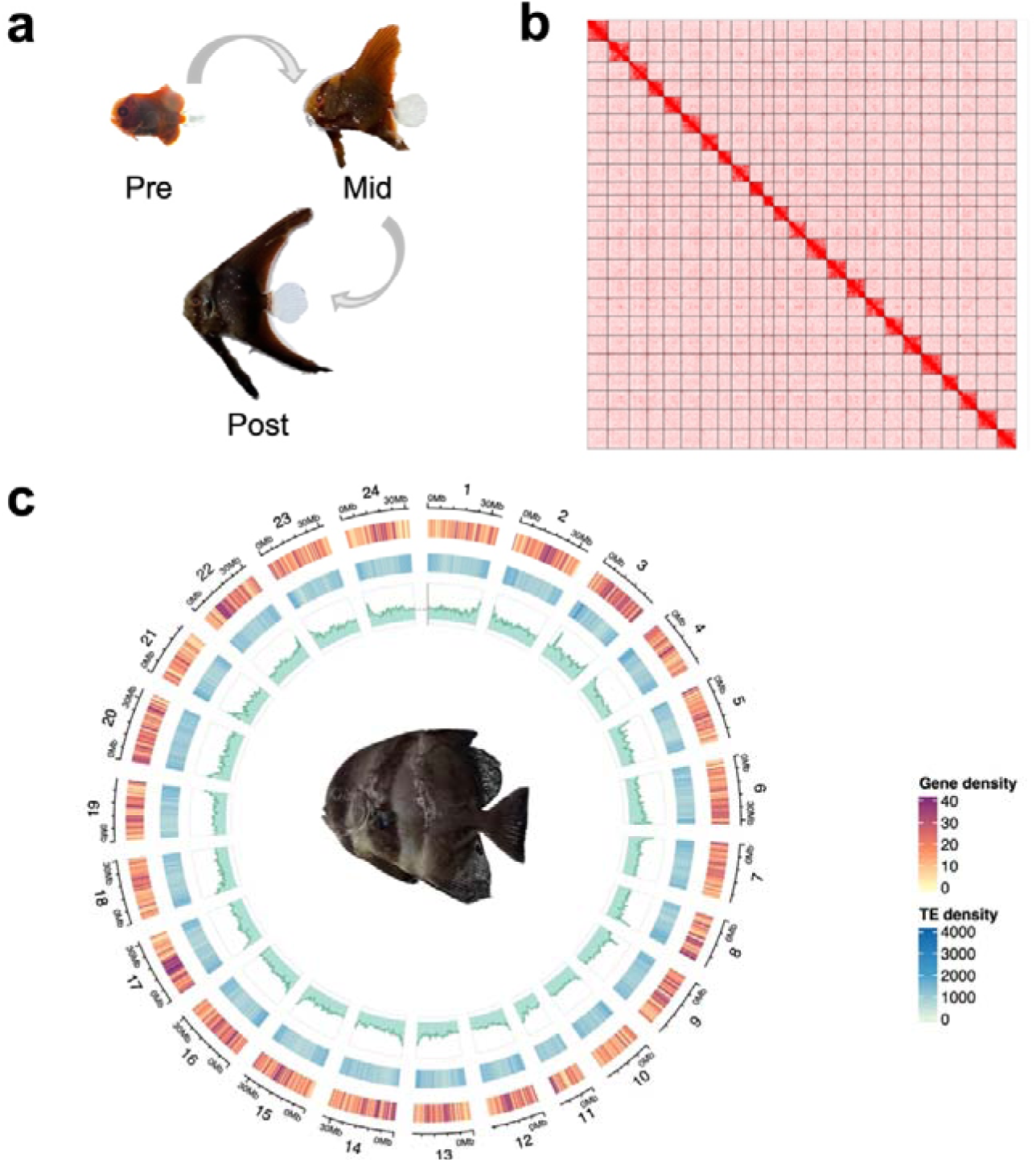
Genomic features of orbicular batfish. **a.** Pre-, mid-, and post-metamorphic stages during the larva-to-juvenile transition for fin sampling. **b.** Hi-C contact map. **c.** Circos plot of orbicular batfish genome assembly. From the inner to the outer track: GC content, transposable element (TE) density, and gene density. A three-year-old orbicular batfish is shown inside the circle.

### 2.3 DNA and RNA extraction and sequencing

The genomic DNA from muscle tissue was isolated using the Monarch Genomic DNA Purification kit (NEB, #T3010) following the manufacturer’s protocol and used for both Illumina short-read and Oxford Nanopore long-read sequencing. DNA quality and integrity were evaluated by NanoDrop spectrophotometry (Thermo Fisher Scientific, USA) and 1.5% agarose gel electrophoresis. For short-read sequencing, genomic DNA was sent to Berry Genomics (Hong Kong, China), where DNA libraries were prepared and sequenced on the Illumina NovaSeq 6000 platform (Illumina Inc., San Diego, CA, USA) in a 150-bp paired-end mode. For long-read sequencing, libraries were generated using the LSK-110 library preparation kit (ONT, Oxford, United Kingdom) according to the standard protocol and sequenced on a MinION platform (ONT) with an R10.4.1 Flow Cell (FLO-MIN114, ONT).

For Hi-C sequencing, sample preparation, extraction, and library preparation were performed according to the restriction enzyme Pore-C (RE-Pore-C) protocol of Oxford Nanopore Technologies, with the MboI restriction enzyme used to digest the genomic DNA. The sequencing was performed by Berry Genomics on the Illumina NovaSeq 6000 platform using 150-bp paired-end reads.

For RNA sequencing, total RNA was extracted from all samples using the TaKaRa MiniBEST Universal RNA Extraction Kit (Takara Bio Inc., Kusatsu, Shiga, Japan). RNA integrity was assessed with an Agilent TapeStation system, and RNA quantity was measured using a NanoDrop spectrophotometer (Thermo Fisher Scientific, USA). RNA samples were sent to Benagen Co., Ltd. (Wuhan, China) for library preparation and sequenced on the DNBSEQ-T7 platform (BGI Inc., Shenzhen, China) in 150-bp paired-end mode.

### 2.3 Genome assembly

To understand the genomic characteristics of orbicular batfish, we employed the *k-mer* frequency distribution analysis to estimate its genome size and genome complexity. The *k-mer* count histogram (*k*LJ=LJ21) was obtained from the Illumina paired-end sequencing data using Jellyfish v2.3.0^19^. Subsequently, the genome size and heterozygosity were estimated by GenomeScope v2.0^20^ after removing the *k-mers* with abnormal depth.

Prior to genome assembly, the raw sequencing data underwent quality control. We utilized fastp v0.23.4^21^ to trim adapters and eliminate low-quality from short reads, while Filtlong v0.3.1 (https://github.com/rrwick/filtlong) was deployed to remove low-quality and overly short Nanopore long reads. The assembly process began with the construction of a draft genome using Flye v2.9.4^22^ with Nanopore reads. To mitigate the error rate associated with Nanopore sequencing, the contigs was initially polished using long reads with Racon v1.4.20^23^, followed by additional polishing using high-accuracy Illumina short reads with Pilon v1.24^24^ to further refine the assembly. The redundant contigs were removed using purge_dups v1.2.6^25^. Hi-C data was utilized to further construct chromosome-level genome assembly of orbicular batfish. The clean Hi-C reads were first mapped to the assembled sequences utilizing BWA v0.7.18-r1243-dirty^26^. Subsequently, Juicer v2.0^27^ was applied to obtain the valid Hi-C contacts with duplicates removed, and 3D-DNA v180922^28^ was used to anchor the assembled contigs onto chromosomes. Finally, Juicebox v2.15^29^ was used to visualize and manually correct the resulting chromosomal scaffolds. The completeness of the final assembly was assessed by BUSCO (Benchmarking Universal Single Copy Orthologs) v5.7.1^30^ analysis, based on 3,640 orthologs from the Actinopterygii_odb10 database.

### 2.4 Genome annotation

For repeat annotation, we first constructed a *de novo* repeat library using RepeatModeler v2.0.5^31^, with RECON v1.0.8^32^ and RepeatScout v1.0.6^33^ to detect repetitive elements in the genome. High-quality, non-redundant long terminal repeat (LTR) retrotransposons were identified by LTR_retriever v2.9.0^34^ and LTRharvest included in GenomeTools suite v1.5.9^35^. In addition, RepeatMasker v4.1.6^36^ was used to identify homolog repetitive elements in the genome based on the lineage-specific repeats from Dfam v3.6^37^ and Repbase^38^. Repetitive sequences identified by both *de novo* and homology-based approaches were then searched against and annotated in the genome using RepeatMasker.

Protein-coding genes were predicted using a combination of *ab initio*, homology-based, and transcript-based approaches. After RNA-seq reads were aligned to the repeat-masked genome using HISAT2 v2.2.1^39^, BRAKER v3.0.8^40^ was employed for *ab initio* gene prediction based on models trained with GeneMark-ETP v1.0^41^ and AUGUSTUS v3.5.0^42^. For homology search-based gene prediction, the protein sequences from six related species, including *Morone saxatilis*, *Gasterosteus aculeatus*, *Larimichthys crocea*, *Oryzias latipes*, *Oreochromis niloticus*, and *Takifugu rubripes*, were aligned to the assembly by exonerate v2.4.0^43^. In addition, RNA-seq data from the liver, kidney, and spleen were assembled into transcripts using Trinity, followed by gene structure prediction with PASA pipeline v2.5.3^44^, which employed BLAT v35.1^45^, GMAP v2024-05-20^46^, and Minimap2 v2.28^47^ to align the assembled transcripts to the genome. To assess the completeness of predicted gene sets, the BUSCO analysis was also performed on the predicted protein sequences using the Actinopterygii_odb10 database. EggNOG-mapper v2.1.12^48^ and InterProScan v5.69-101.0^49^ were used for gene function annotation and Gene Ontology (GO) annotation.

### 2.5 Gene family analysis

Fourteen representative species were selected to perform the phylogenetic analysis with orbicular batfish, including large yellow croaker (*Larimichthys crocea*), striped bass (*Morone saxatilis*), copperband butterflyfish (*Chelmon rostratus*), Japanese pufferfish (*Takifugu rubripes*), humphead wrasse (*Crassilabrus undulatus*), three-spined stickleback (*Gasterosteus aculeatus*), ocellaris clownfish (*Amphiprion ocellaris*), Nile tilapia (*Oreochromis niloticus*), medaka (*Oryzias latipes*), tongue sole (*Cynoglossus semilaevis*), tiger tail seahorse (*Hippocampus comes*), Atlantic cod (*Gadus morhua)*, zebrafish (*Danio rerio*), and Asian arowana (*Scleropages formosus)*. Protein sequences from the 15 species were clustered using OrthoFinder v2.5.5^50^ with default parameters. The protein sequences of single-copy orthologs were aligned with MUSCLE v5.1.l^51^ and the corresponding coding sequence alignments were generated using pal2nal v14 based on the protein alignments. Low-quality aligned coding sequences were then removed using Gblocks v0.91b^52^ with the parameter “-t=c”. A concatenated matrix was generated from all filtered alignments, which was used to construct a maximum-likelihood tree using IQ-TREE v2.3.5^53^ with 1,000 bootstrap replicates. The optimal nucleotide substitution model was determined by ModelFinder^54^ based on the Bayesian information criterion. The divergence times between individual species were estimated using MCMCtree in the PAML software v4.10.7^55^, based on 4D site alignments and four calibration time points (large yellow croaker–medaka: ∼100–130LJMya, large yellow croaker–Atlantic cod: ∼128–165 Mya, large yellow croaker–zebrafish: ∼180–251.5 Mya, and large yellow croaker–Asian arowana: ∼215-297.9 Mya) on the TIMETREE website (https://timetree.org/).

To investigate the gene families evolution in orbicular batfish, CAFE5^56^ was employed to estimate gene families expansion and contraction in the constructed species tree, with the estimated divergence times between species as input. The clusterProfiler package v4.10.0^57^ in R was used to perform the GO enrichment analysis of gene families expanded and contracted in orbicular batfish.

### 2.6 Identification of positively selected gene

The selection pressure on orbicular batfish was analyzed using the CODEML program in PAML software v4.10.7. To identify positively selected genes (PSGs), we utilized the branch-site model, where the null hypothesis assumed that the Ka/Ks ratio for each site on every branch was 1, while the alternative hypothesis allowed certain sites on the foreground branch showing *Ka/Ks* ratios > 1. The Chi-square test was utilized to calculate the *P*-values based on the likelihood ratio test (LRT). Genes with a False Discovery Rate (FDR)-adjusted *P*-value < 0.05 and containing sites with a posterior probability > 0.95 in the Bayes Empirical Bayes (BEB) analysis were considered to be under positive selection.

### 2.7 Differential gene expression analysis

Raw RNA-seq reads from mixed dorsal, pelvic, and anal fins at different developmental stages were first filtered using fastp v0.23.4, and the cleaned reads were then mapped to the genome using STAR v2.7.11b^58^. Gene expression levels were quantified with RSEM v1.3.3^59^. Prior to differential expression analysis, lowly expressed genes were removed, and only genes with read counts ≥10 in at least three samples were retained. Genes that were significantly differentially expressed between any two developmental stages (adjusted *p*-value < 0.05, ∣log2FoldChange∣ > 1) were identified using DESeq2 v1.42.1^60^. Since one sample from the pre-metamorphic stage was extracted in a different batch from the others, batch was included as a covariate in the DESeq2 model. All significantly differentially expressed genes were clustered using the Mfuzz package v2.62.0^61^ in R. Batch effects were further removed in the expression matrix using removeBatchEffect function from the limma package v3.58.1^62^. The optimal number of clusters was determined based on the minimum centroid distance method using the Dmin function in Mfuzz v2.62.0, as well as the elbow method of Within-Cluster Sum of Squares. Genes with membership values greater than 0.3 in each cluster were defined as high-confidence cluster members and were subjected to GO enrichment analysis using the clusterProfiler package v4.10.0^63^.

### 2.8 Identification of *hox* gene clusters

We identified *hox* genes in orbicular batfish, copperband butterflyfish, and large yellow croaker by performing BLASTP v2.15.0^64^ searches using 49 zebrafish Hox proteins as queries, with a cutoff of e-value < 1×10^−20^. HMMER v3.4^65^ was then used to confirm the presence of the homeodomain (Pfam accession: PF00046) in the identified Hox proteins. To further validate the *hox* genes, we constructed a protein phylogenetic tree of hox genes from orbicular batfish, copperband butterflyfish, large yellow croaker, Japanese pufferfish, and zebrafish using IQ-TREE v2.3.5. The phylogenetic tree was visualized with iTOL v7.6^66^.

## 3. Results

### 3.1 High-quality genome assembly of orbicular batfish

Nanopore long-read sequencing generated 2.38 million reads with an average read length of 12.93 kb, yielding 30.82 Gb of total bases (Supplementary Table 1). In parallel, short-read sequencing produced 697.16 million reads, corresponding to 104.57 Gb of total bases, while Hi-C sequencing generated 36.97 Gb total bases. These datasets achieved approximately 43×, 146×, and 52× coverage of the genome, respectively.

Based on k-mer analysis of the short-read data, the genome size was estimated to be 655.47 Mb, with a heterozygosity rate of 0.249% (Supplementary Fig. S1). The final genome assembly shows a total size of 715.44 Mb, with a total of 280 scaffolds (Fig. 1b and Supplementary Table 2). The contig N50 of the assembly is 13.27 Mb and the scaffold N50 is 31.08 Mb. Using Hi-C data, the assembled sequences were further anchored onto 24 chromosomes (Fig. 1b), with chromosome sizes ranging from 16.43 Mb to 34.87 Mb. Assessment of assembly completeness using BUSCO revealed 99.5% complete BUSCOs (Supplementary Table 2), indicating that the genome is highly complete. Furthermore, mapping the high-accuracy short reads back to the assembled genome showed that 99.90% of the reads were successfully aligned, covering 99.93% of the assembly, which further supported the high accuracy and reliability of the genome assembly.

For repeats prediction, by combining homology-based and *de novo* methodologies based on RepeatModeler and RepeatMasker, we estimated that approximately 26.61% of the genome consisted of repetitive sequences, corresponding to 190.36 Mb (Fig. 1c and Supplementary Table 3). For protein-coding gene prediction, our approach, combining the *ab initio* prediction, transcript-based strategies, and protein alignment, predicted a total of 25,544 protein-coding genes and 243,568 exons within our genome assembly (Fig. 1c and Supplementary Table 4). These genes exhibited an average gene length of 12,420 bp and an average exon length of 165 bp. The BUSCO score for the predicted genes is 94.4% (Supplementary Table 4), indicating a high level of completeness in the predictions. For gene functional annotation, eggNOG-mapper and InterProScan yielded 22,999 (90.03%) and 23,769 (93.05%) hits, respectively. After integrating the results from both annotation methods, a total of 24,018 (94.02%) genes were successfully assigned with putative functional.

### 3.2 Gene family analysis

To elucidate the evolutionary history of orbicular batfish, we constructed a phylogenetic tree incorporating 14 other representative teleost species. Large yellow croaker, striped bass, and copperband butterflyfish were selected due to their presumed close phylogenetic affinities, while the remaining species were included to provide a cover a broad range of teleosts. Our analysis revealed that the orbicular batfish and copperband butterflyfish cluster together (Fig. 2a and Supplementary Fig. S2), strongly supporting a close evolutionary relationship that aligns with their shared morphological traits. The divergence time between these two species was estimated to be approximately 73.25 million years ago (Mya).

**Figure 2.**
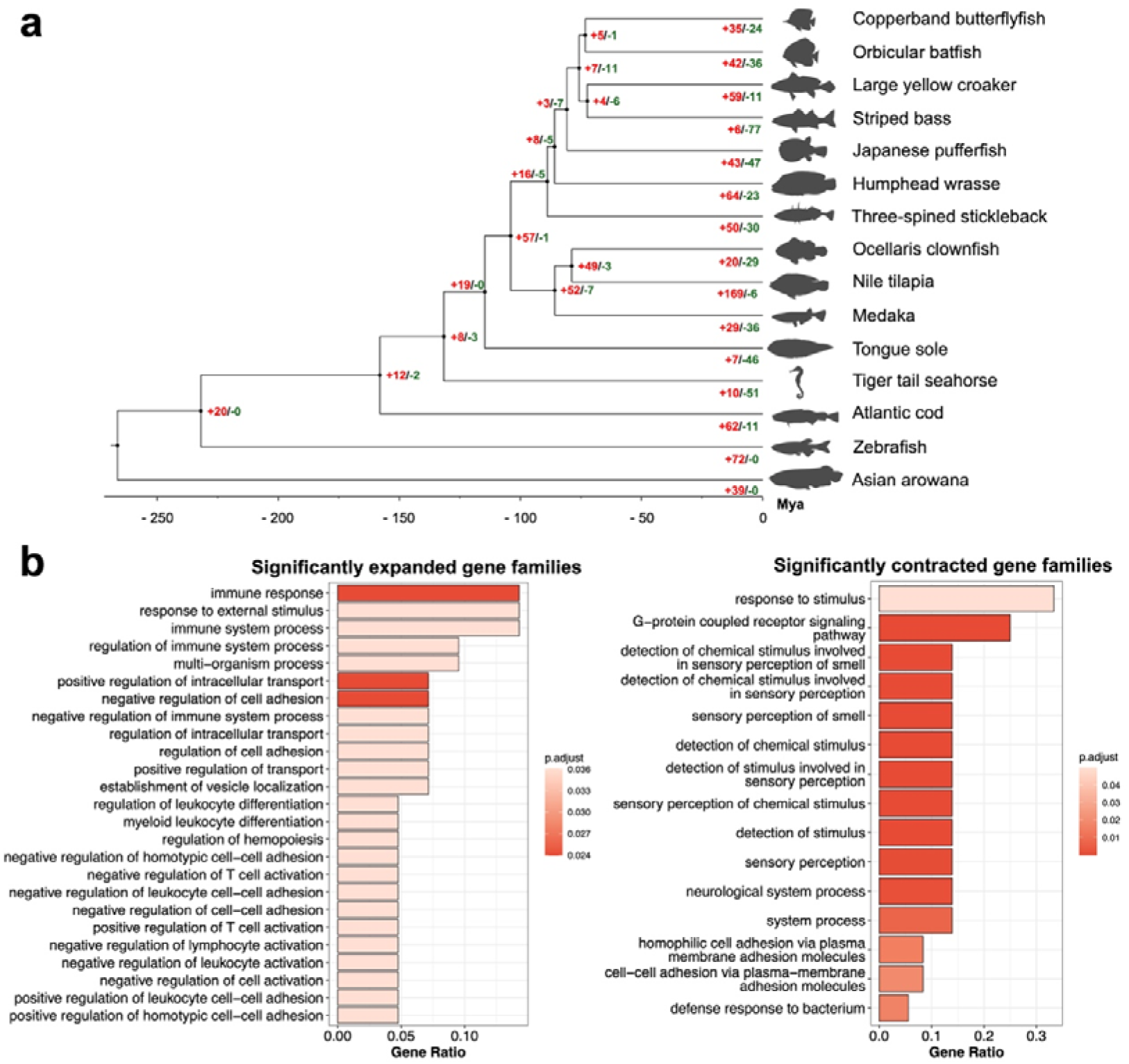
Gene family evolution in orbicular batfish. **a.** Time-calibrated phylogenetic tree of orbicular batfish and 14 other representative fish species. Red and green numbers beside each node indicate the numbers of significantly expanded and contracted gene families, respectively. **b.** Top 25 significantly enriched GO biological process (BP) terms for significantly expanded and contracted gene families in orbicular batfish.

To uncover the genetic basis underlying the distinctive morphological development of orbicular batfish, we further investigated evolutionary dynamics at the gene family level. Ortholog clustering across the 15 genomes identified a total of 21,657 gene families, with 18,437 present in orbicular batfish. Compared to its most recent common ancestor shared with copperband butterflyfish, orbicular batfish genome exhibited 511 expanded and 1,051 contracted gene families (Supplementary Fig. S3a), among which 42 and 36 families underwent significant expansion and contraction, respectively (Fig. 2a). GO enrichment analysis of the 42 significantly expanded gene families revealed that they were predominantly enriched in immune-related pathways (BH-adjusted *p*-value < 0.05), such as immune response, immune system process, and regulation of T cell activation (Figure. 2b). Conversely, the 36 significantly contracted families were primarily enriched in sensory and stimulus-response pathways, such as sensory perception of smell and detection of chemical stimulus (Figure 2b). These genomic changes likely reflect the accumulation of adaptive potential in response to the ecological niche shifts and environment challenges associated with metamorphosis in this species. Interestingly, broadening the analysis to include all 511 expanded gene families revealed a distinct functional pattern. This comprehensive set was significantly enriched in fundamental developmental processes, evidenced by terms such as anatomical structure development, system development, and animal organ development (Supplementary Fig. S3b). These findings suggest that orbicular batfish genome maintains a robust and broad genetic foundation to support its unique morphological transformations. In contrast, no significantly enriched GO terms were detected among all contracted gene families.

### 3.3 Gene expression analysis during metamorphosis

Genome-level evolutionary analysis described the genetic toolkit of orbicular batfish. To further reveal the regulatory networks of metamorphosis, we explored the dynamic transcriptomic changes across three developmental stages spanning the transition from larva to juvenile, representing the pre-, mid-, and post-metamorphic stages. We performed RNA sequencing specifically on dorsal, pelvic, and anal fins showing the most pronounced morphological changes to explore the downstream regulatory pathways. On average, 87.88 million reads were obtained for each pooled or individual sample (Supplementary Table 1). Principal component analysis (PCA) of gene expression showed that samples from the three stages were clearly separated along the first and second principal components (Fig. 3a).

**Figure 3.**
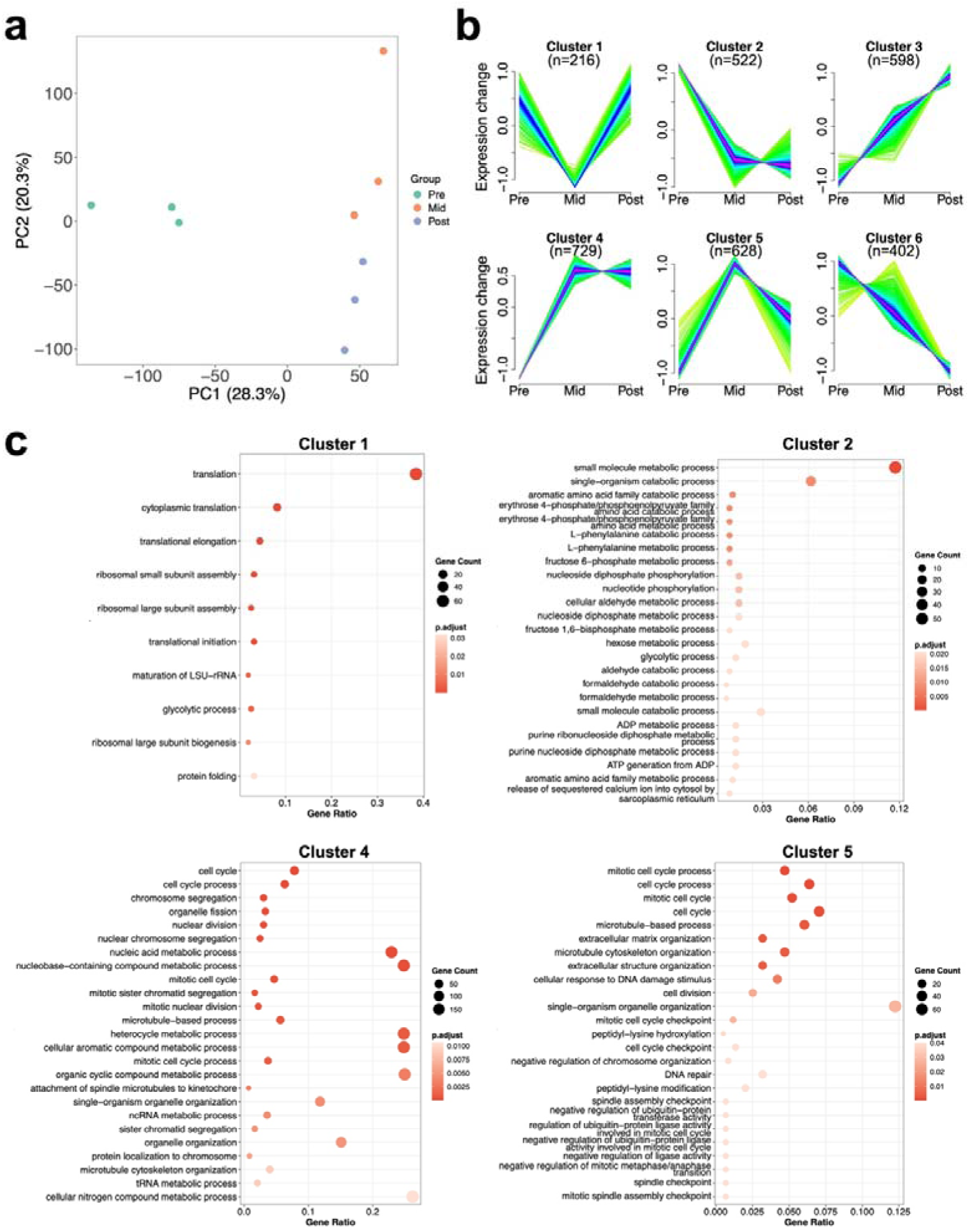
Transcriptomic analysis of dorsal, pelvic, and anal fins during larva-to-juvenile metamorphosis in orbicular batfish. **a.** Principal component analysis (PCA) of fin gene expression at the pre-, mid-, and post-metamorphic stages. **b.** Expression pattern clustering analysis using Mfuzz. The number of genes in each cluster is shown above the corresponding panel. Gene expression data were normalized to a mean of 0 and a standard deviation of 1. **c.** Top 25 significantly enriched GO biological process terms for each cluster. No significantly enriched GO terms were identified for clusters 3 and 6.

Differential expression analysis across the pre-, mid-, and post-metamorphic stages identified a total of 3,095 genes that were significantly altered in at least one pairwise comparison between stages. To further characterize gene expression dynamics during metamorphosis, we applied Mfuzz to group genes with similar expression patterns across all stages in order to identify core gene modules specifically activated at particular stages. The optimal clustering result was obtained when the number of clusters was set to six, and each of the 6 clusters exhibited a distinct temporal expression pattern (Fig. 3b and Supplementary Fig. S4). Among these clusters, cluster 4 (729 genes) and cluster 5 (628 genes) contained the largest numbers of genes and were both rapidly activated at the mid-metamorphic stage. Using high-confidence genes in each cluster for enrichment analysis, we found that both clusters were significantly enriched in cell cycle–related pathways, such as mitotic cell cycle, chromosome segregation, and spindle assembly checkpoint, indicating an activation of cell proliferation during mid-metamorphosis (Fig. 3c). Interestingly, while the expression of cluster 5 declined at the post-metamorphic stage, cluster 4 maintained a relatively stable high expression level, suggesting that part of the cell proliferation-associated regulatory program persists until the post-metamorphic stage. In contrast to the activation of cell proliferation-related pathways, some pathways were suppressed during mid-metamorphosis. Cluster 2 (522 genes), which was predominantly enriched in fundamental metabolic pathways, showed a marked downregulation at the mid-metamorphic stage and remained at a relatively low expression level thereafter (Fig. 3b). Specifically, these metabolic genes were broadly involved in energy and carbohydrate metabolism, including fructose 6-phosphate metabolic process, ATP generation from ADP, and glycolytic process, as well as amino acid metabolism, such as aromatic amino acid family catabolic process and L-phenylalanine metabolic process (Fig. 3c). Cluster 1 (216 genes), which displayed a transient decrease at the mid-metamorphic stage and thus showed an inverted V-shaped trend, was mainly enriched in translation-related pathways and was also enriched in glycolytic process (Fig. 3b and c). These expression patterns may suggest a reallocation of energy and resources during metamorphosis, where the temporary suppression of metabolism and translation may help activate the increased demand for cell division, supporting the elongation of fins. In addition, cluster 3 (598 genes) and cluster 6 (402 genes) showed continuously increasing and decreasing expression trends, respectively, but did not exhibit significant enrichment for any GO terms, possibly because these genes are functionally diverse.

### 3.4 Positively selective genes associated with fin remodeling

To further understand the genetic basis of adaptive morphological innovation in orbicular batfish, we examined signatures of positive selection across the genome. In total, we identified 300 significant positively selected genes (PSGs). Functional annotation of these PSGs revealed a broad selection within genes involved in extracellular matrix (ECM) organization, such as *col18a1*, *col21a1*, *matn1*, and *sgcb*, as well as genes governing cell cycle and DNA repair, such as *e2f1*, *brca2*, *xrcc6*, *nhej1*, *cdkn1c*, and *mcph1* (Supplementary Table 5).

To uncover the evolutionary targets potentially associated with local fin remodeling, we intersected the PSGs with the differentially expressed genes identified specifically from fin transcriptomes during metamorphosis. This analysis highlights a set of candidate genes involved in cell cycle regulation, skeletal development, and ECM remodeling (Fig. 4). Among them, two genes related to cell cycle and DNA repair, *ankfn1* and *mcph1*, were significantly activated at the mid-metamorphotic stage. In skeletal regulation, two PSGs, *cyp24a1* and *sema4d*, showed relatively high expressed exclusively at the pre-metamorphic stage, followed by a significant downregulation at the post-metamorphic stage. Furthermore, *sdc4* and *serpine1*, two PSGs involved in ECM remodeling, also showed significant expression changes. *Sdc4* exhibited higher expression at the pre-metamorphic stage and was significantly downregulated at the post-metamorphic stage, while *serpine1* was significantly upregulated during the mid-metamorphic stage and maintained high-level expression through the post-metamorphic stage.

**Figure 4.**
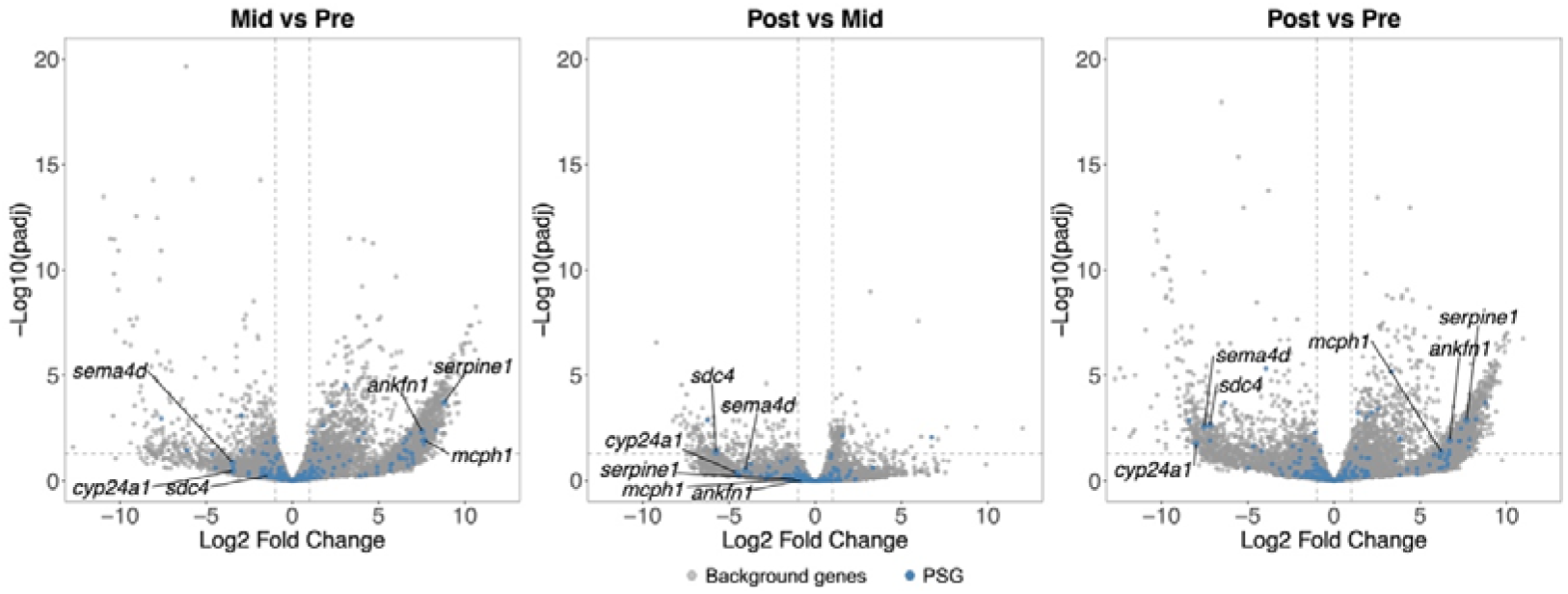
Expression changes of PSGs in the dorsal, pelvic, and anal fins across the pre-, mid-, and post-metamorphic stages. Blue dots represents PSGs, while gray dots represent other genes.

### 3.5 *Hox* gene expression dynamics

*Hox* genes are important developmental regulators that play essential roles in vertebrate morphogenesis and organogenesis. Because of whole-genome duplication and gene loss during evolution, the composition of *hox* clusters varies substantially among vertebrate lineages. We identified 49 *hox* genes in orbicular batfish and each Hox protein was clustered with its orthologs from copperband butterflyfish, large yellow croaker, butterflyfish, Japanese pufferfish, and zebrafish (Fig. 5a and Supplementary Fig. S5). The *hox* clusters were conserved between orbicular batfish and copperband butterflyfish. Compared with Japanese pufferfish, however, orbicular batfish retains four additional genes, *hoxa7a*, *hoxb7a*, *hoxc1a*, and *hoxc3a*.

**Figure 5.**
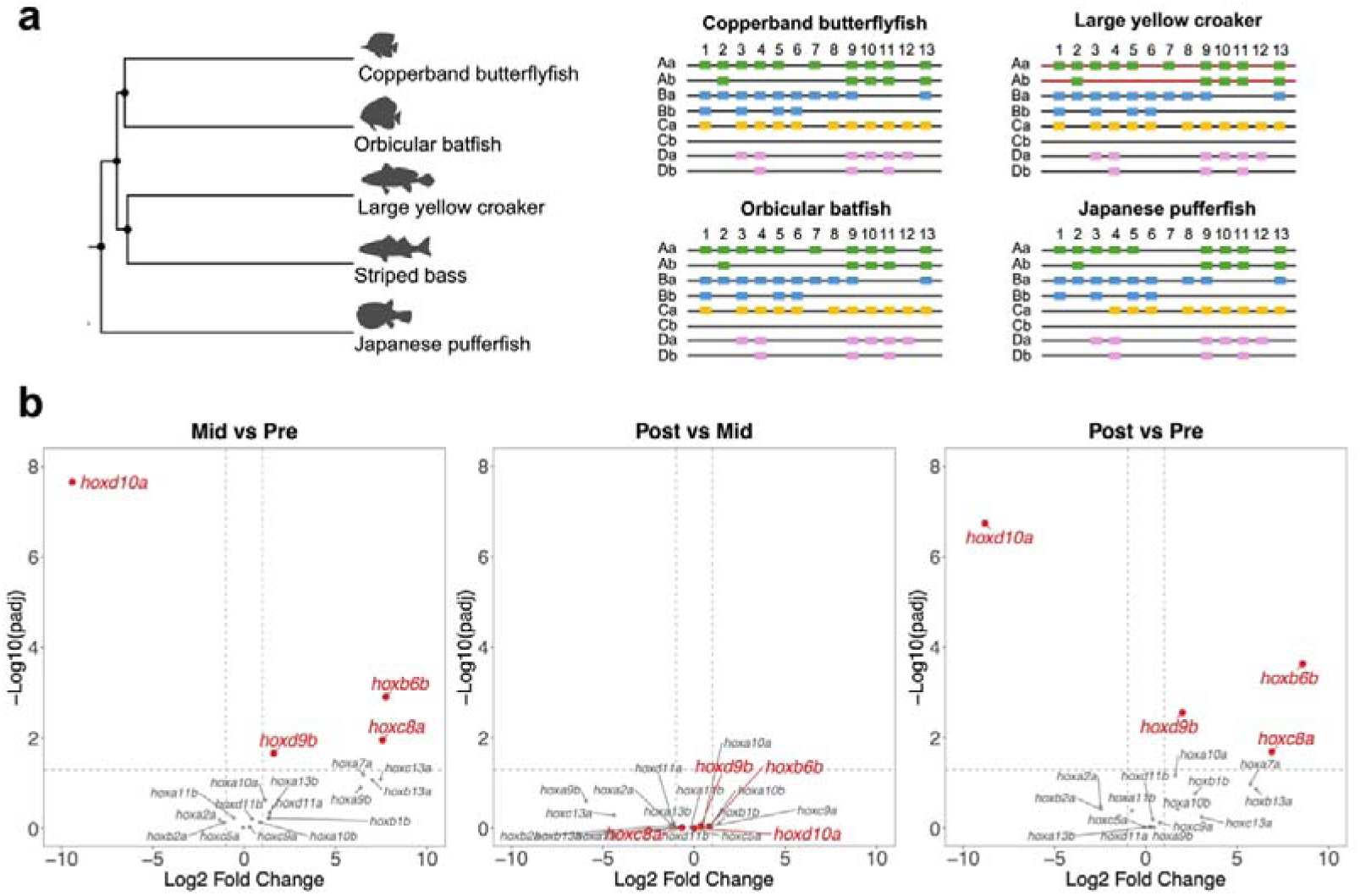
Expression changes of *hox* genes across the pre-, mid-, and post-metamorphic stages. **a.** Identification of *hox* genes. **b.** Volcano plots showing *hox* gene expression in the dorsal, pelvic, and anal fins of orbicular batfish during metamorphosis. Lowly expressed *hox* genes were excluded, and only genes with read counts of at least 10 in at least three samples are shown.

We further examined the expression dynamics of *hox* genes in dorsal, pelvic, and anal fins during metamorphosis in orbicular batfish. Notably, *hoxb6b*, *hoxc8a*, *hoxd9b*, and *hoxd10a* showed significant expression changes at the mid-metamorphic stage (Fig. 5b). Among them, *hoxd10a* was downregulated, whereas *hoxb6b, hoxc8a*, and *hoxd9b* were upregulated, and their expression levels remained relatively stable from the mid-to post-metamorphic stages. *Hoxd10a* has previously been reported to be associated with dorsal fin development in zebrafish^67^. In stickleback, the *hoxdb* locus, including *hoxd9b*, has been linked to spine elongation and spine number^68^.

## 4. Discussion

Metamorphosis in teleost fishes is a life-history transition that bridges ecological shifts with coordinated morphological, physiological, and behavioral adaptations. Orbicular batfish (*Platax obicularis*) serves as a representative example of extreme morphological innovation, characterized by accelerated growth of its fins relative to the body during the larval-to-juvenile transition. While previous studies on fish metamorphosis have predominantly relied on single-dimensional transcriptomic profiling, our study provides a comprehensive evolutionary developmental perspective. By assembling a high-quality chromosome-level genome and integrating evolutionary genomic signatures with spatiotemporal transcriptomic dynamics, we uncover the genetic architecture underlying this extreme trait. Our findings demonstrate that the metamorphosis of orbicular batfish is not merely a localized developmental event, but a systemic transition driven by the convergence of genome-level adaptive evolution and transcriptional reprogramming.

### 4.1 Refinement of genomic repertoire provides the potential for ecological adaptation and development

Expansion and contraction of gene families essentially reflect genomic adjustments to environmental pressures over evolutionary timescales. In orbicular batfish, metamorphosis is accompanied by a marked ecological transition from a pelagic larval lifestyle in the open water to a juvenile habitat associated with coral reef environments. This habitat shift exposes individuals to substantially different biotic and abiotic conditions, thereby imposing considerable environmental challenges. Previous studies have shown that fish generally exhibit environmental plasticity, enabling them to cope with external changes encountered during development or across life-history stages^69,70^. The long-term maintenance of such plasticity is thought to be achieved through epigenetic regulation or evolutionary adaptation at the genomic level^69^. In this study, we identified significant expansion of immune-related gene families and significant contraction of sensory-related gene families in orbicular batfish. This pattern of gene family expansion and contraction may represent genomic remodeling associated with adaptation to the environmental pressures accompanying metamorphosis. Environmental changes often have direct impacts on the immune system, as fishes must respond to multiple stressors in different habitats, including pathogen exposure, temperature, oxygen, salinity, and pH changes^69,71^. Therefore, the expansion of immune-related gene families may contribute to improved survival in new environments. At the same time, sensory systems are essential for fishes to function in specific habitats. During habitat transition, fishes often need to recalibrate how they perceive environmental information and stimuli in multiple sensory ways, including chemosense, vision, and magnetoreception^72,73^. Therefore, the contraction of sensory-related gene families may reflect the adjustment of sensory system in response to the ecological demands of the new habitat. In addition, analysis of all expanded gene families revealed that they were significantly enriched in development-related pathways. This pattern suggests that extensive gene family expansion during the evolutionary history of orbicular batfish may have provided the necessary genetic redundancy and broad developmental toolkit that supports its complex morphological transformations.

### 4.2 Energy and resource trade-offs during fin remodeling

Ecological adaptation during metamorphosis requires not only long-term evolutionary remodeling, but also short-term coordinated developmental regulation. Since energy and resources are inherently limited, the enhancement of specific biological functions often comes at the expense of others, and such trade-offs are widespread across different taxa^74,75^. For example, juvenile brown stingrays under food restriction exhibit a trade-off between metabolic maintenance and growth ^76^, while juvenile rainbow trout in winter environment balance metabolic and growth rates to enhance survival^77^. The extreme morphological remodeling during orbicular batfish metamorphosis may represent a similar biological trade-off. During the middle stage of fin metamorphosis, we observed suppression of basal metabolism and translation processes, accompanied by strong activation of cell cycle pathways. This expression pattern suggests that orbicular batfish may employ a developmental strategy that prioritizes rapid local growth during the critical metamorphic period. Cell proliferation is the essential process for fin development and regeneration^78^, while the concurrent downregulation of carbohydrate metabolism, amino acid metabolism, and certain non-essential protein synthesis may effectively redirect available energy and resources toward targeted fin remodeling. This reprogramming indicates that metamorphosis in orbicular batfish is not simply a uniform physiological shift, but rather a fine-tuned reprioritization of biological processes to balance organismal survival and morphological innovation. Notably, similar patterns of metabolic reprogramming have also been reported during metamorphosis in amphibians and insects^79,80^, suggesting that the temporary reconfiguration of basal physiological functions to ensure critical developmental events may represent a common strategy underlying dramatic metamorphic remodeling. Furthermore, we detected differential expression of several hox genes, including *hoxb6b*, *hoxc8a*, *hoxd9b*, and *hoxd10a*, in metamorphosing fins. Given the conserved role of *hox* genes in axial patterning and appendage development of vertebrates^81^, these expression changes suggest a potential contribution to the allometric elongation of the dorsal, pelvic, and anal fins. However, their precise roles in this process remain to be functionally validated.

### 4.3 Adaptive evolution of key fin remodeling networks

We have identified a large number of positively selected genes in orbicular batfish, however, an important challenge is to determine which of these selective signals are most relevant to fin remodeling. To address this, we integrated signatures of positive selection with metamorphic expression dynamics. This approach is informative because adaptive evolution can operate across multiple genomic dimensions, including modifying protein-coding sequences and reshaping gene regulation^82,83^. Positive selection on coding sequences may alter the properties or functions of proteins, whereas divergence in gene regulation often drives gene expression shifts^84–86^. Genes bearing both signals are therefore especially compelling candidates, as they may reflect coordinated evolutionary change at both functional and regulatory levels.

Our integrated analysis highlighted three key processes intrinsically linked to fin transformation, including cell proliferation and DNA repair (*ankfn1* and *mcph1*), skeletal regulation (*cyp24a1* and *sema4d*), and ECM remodeling (*sdc4* and *serpine1*). For example, *ankfn1*, predicted to function in mitotic spindle orientation and bipolar cell polarity, and *mcph1*, related to activation of checkpoint and DNA repair^87^, showed both positive selection and mid-metamorphic upregulation. Because cell proliferation is essential for fin development and regeneration, and proliferating cells are particularly vulnerable to DNA damage^78,88^, the coordinated enhancement of these genes may be critical for supporting rapid fin growth during metamorphosis. Likewise, the ECM-related genes *sdc4* and *serpine1* also emerged as strong candidates, as both showed significant expression changes and signatures of positive selection. ECM program is thought to be essential for fin regeneration in zebrafish^89,90^. SDC4 links the ECM to cytoskeletal signaling and is important for tissue regeneration, while SERPINE1 functions as an inhibitor of ECM degradation^91–93^. Furthermore, we observed both selective signatures and post-metamorphic downregulation in *cyp24a1* and *sema4d* related to skeletal regulation. *Cyp24a1* is involved in vitamin D metabolism and calcium homeostasis, and *sema4d* has been reported to inhibit osteoblast differentiation^94,95^. In fish, fin length is generally thought to occur through the distal addition of bony segments^96^. The early-stage specific high expression and selection signatures further supports the role of ossification in excessive fin elongation in orbicular batfish. Together, the convergence of positive selection and stage-specific expression in these pathways suggests that the extreme fin phenotype of orbicular batfish may evolve through a fine-tuned, dual-level adaption of key remodeling networks, bridging regulatory reprogramming with protein-coding refinement.

## 5. Conclusion

Overall, this study provides a high-quality genomic resource for orbicular batfish and investigates the evolutionary and molecular basis of its remarkable metamorphosis. Our findings suggest that extreme fin elongation in this species is not driven by a single developmental change, but instead emerges from the integration of ecological niche-associated genomic adaptation, reprogramming of energy and resource allocation during fin remodeling, and adaptive evolution of key developmental networks. In this framework, metamorphosis in orbicular batfish can be understood as a coordinated process linking environmental transition, physiological reprioritization, and morphological innovation. By combining comparative genomic signals with tissue-specific developmental dynamics, our study not only offers insights into the basis of this distinctive phenotype, but also expands our understanding of ecological adaptation and metamorphosis novelty in teleosts.

## Author contributions

**Yuxuan Zhang:** conceptualization, methodology, formal analysis, investigation, visualization, resources, writing – original draft, writing – review & editing. **Yanwen Shao:** resources, data curation, visualization, writing – review & editing. **Liang Zhong:** resources, data curation, writing – review & editing. **Runsheng Li:** conceptualization, methodology, data curation, supervision, project administration. **Wenlong Cai**: conceptualization, methodology, data curation, supervision, project administration, funding acquisition.

## Acknowledgments

We thank Dr. Purcia Yuen and Dr. Terence W.K. Chung from Aquaculture Fisheries Division (Mariculture Development Section) of Agriculture, Fisheries and Conservation Department in Hong Kong for providing the batfish and assistance with sample preparation and fish transportation.

## Data availability

Raw sequencing data were deposited in the NCBI Sequence Read Archive database under BioProject number PRJNA1477746. The genome assembly has been deposited at the NCBI GenBank under the accession JBZGVX000000000.

