## Supplementary Fig. for "From pelagic to reef: Genomic basis of morphological adaption during life-history transition in the orbicular batfish (*Platax orbicularis*)"

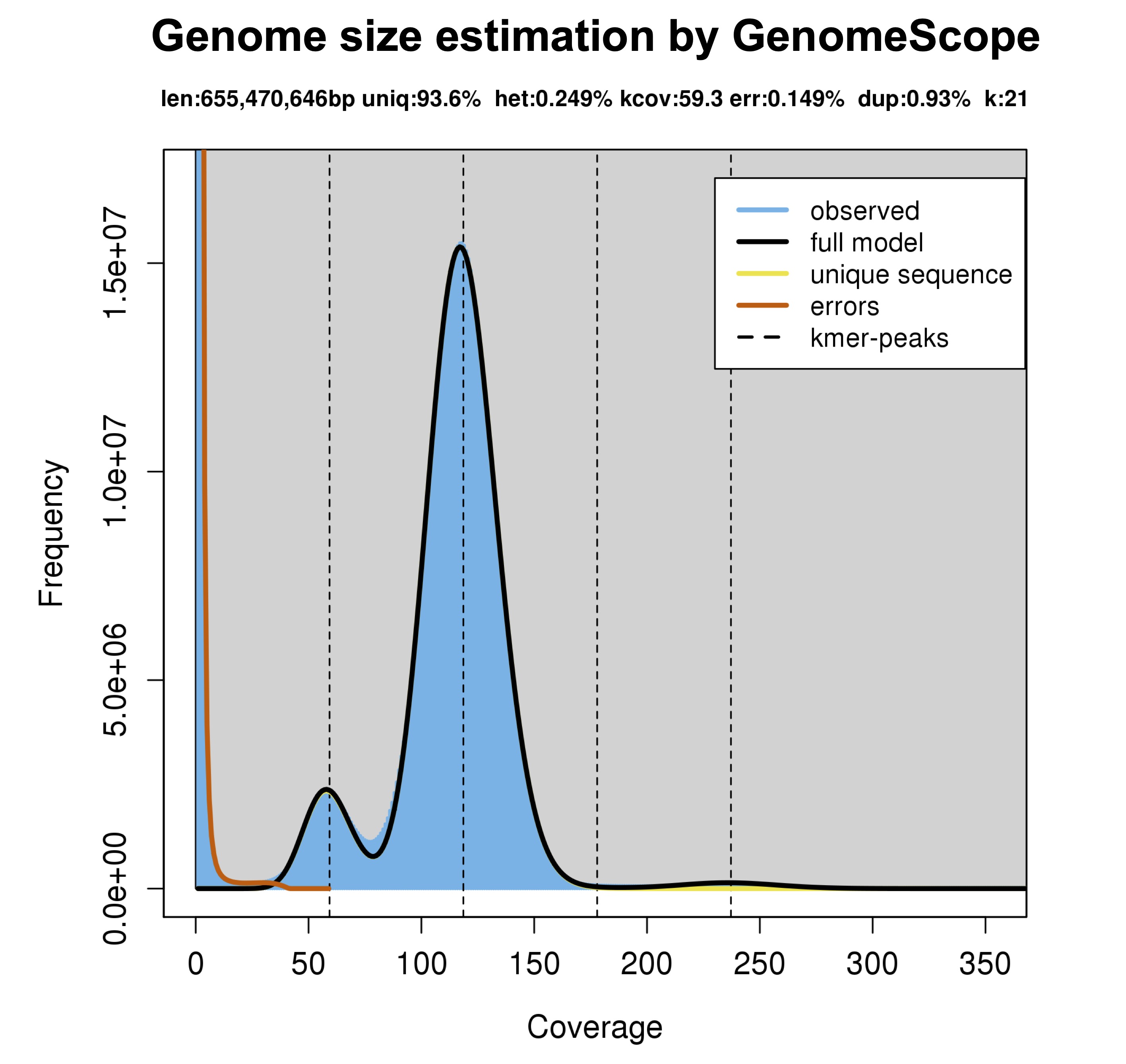

**Figure S1.** Genome size estimation by *k-mer* frequency distribution analysis.

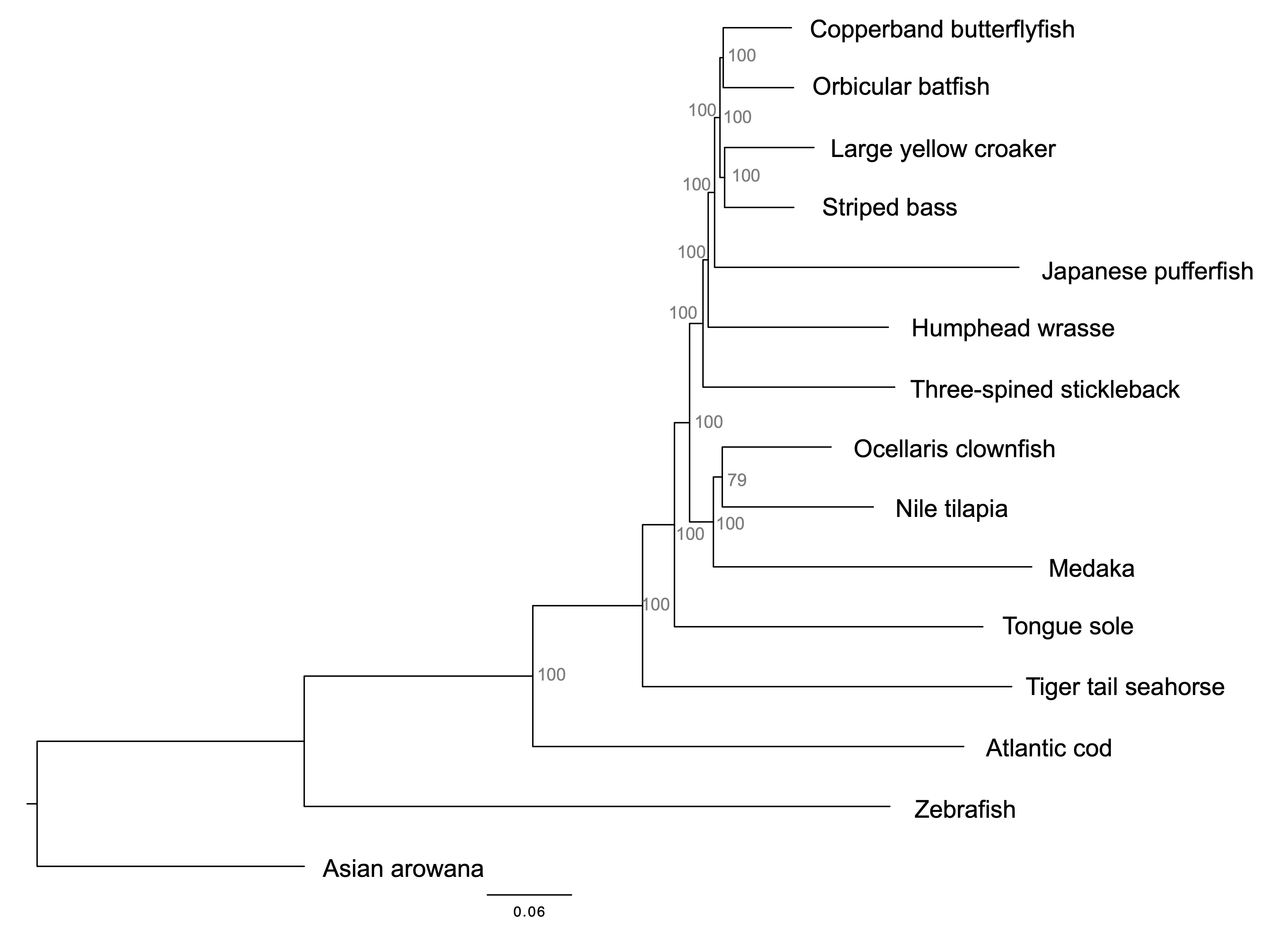

**Figure S2.** Maximum-likelihood phylogenetic tree of orbicular batfish and 14 representative fish species. Numbers at the nodes indicate bootstrap values.

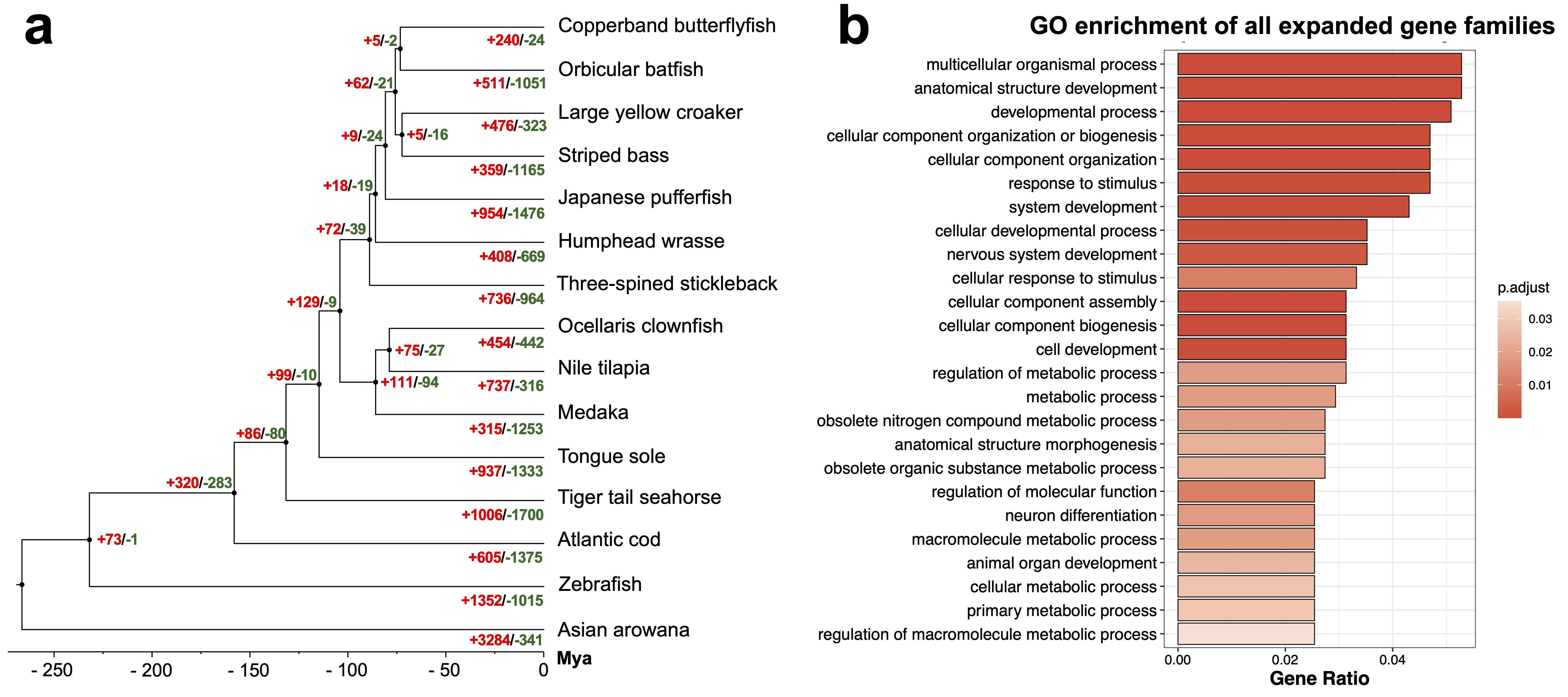

**Figure S3.** All expanded and contracted gene families. **a.** Time-calibrated tree, with the red numbers beside each node indicating the numbers of all expanded gene families and the blue numbers indicating the numbers of all contracted gene families. **b.** Top 25 significantly enriched GO biological process (BP) terms for all expanded gene families in orbicular batfish. No significant GO term was detected for all contracted gene families.

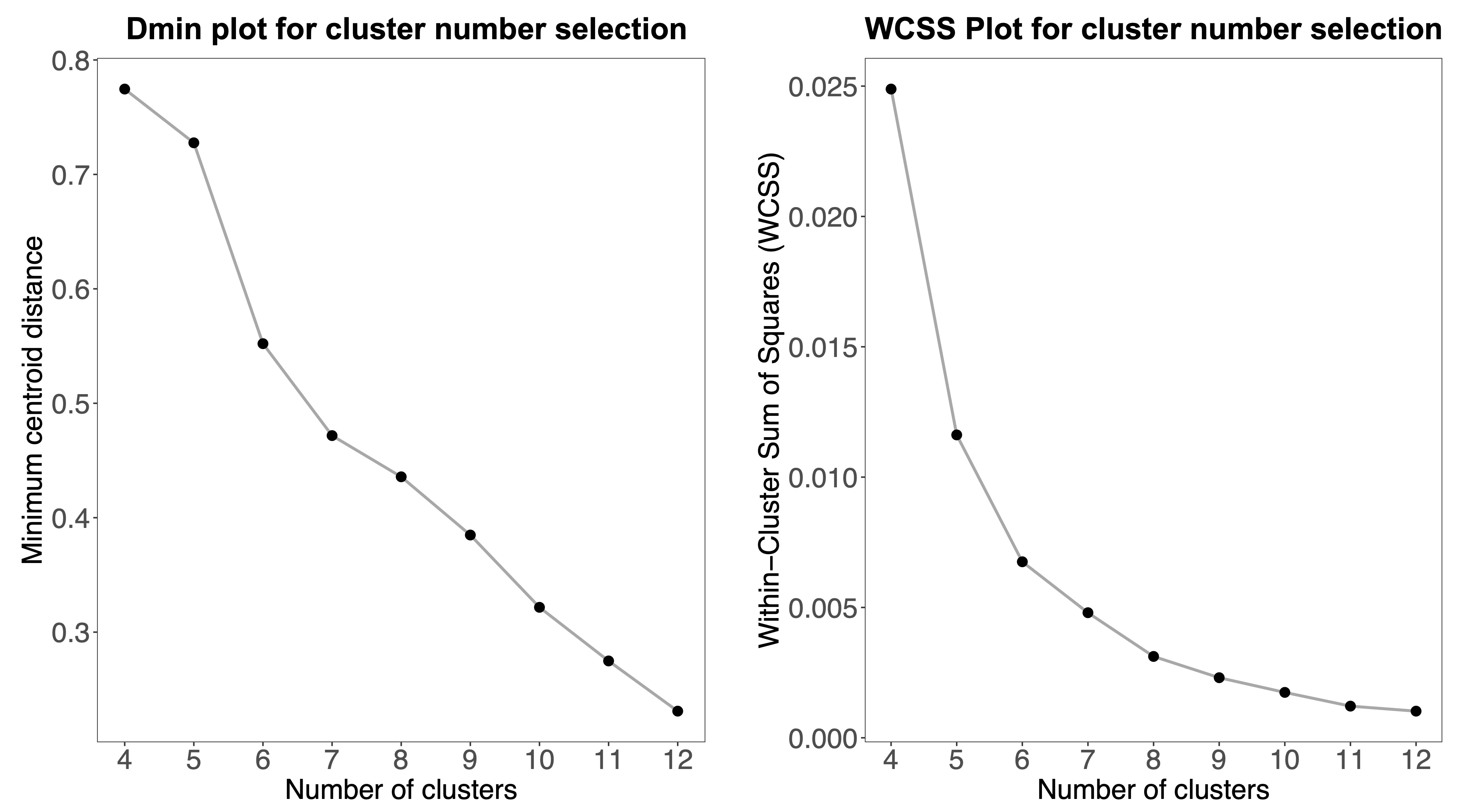

**Figure S4.** Optimal cluster number determination using the minimum centroid distance **(a)** and Within-Cluster Sum of Squares (WCSS) **(b)** methods. Both the minimum centroid distance and WCSS showed a clear elbow point when the cluster number is 6.

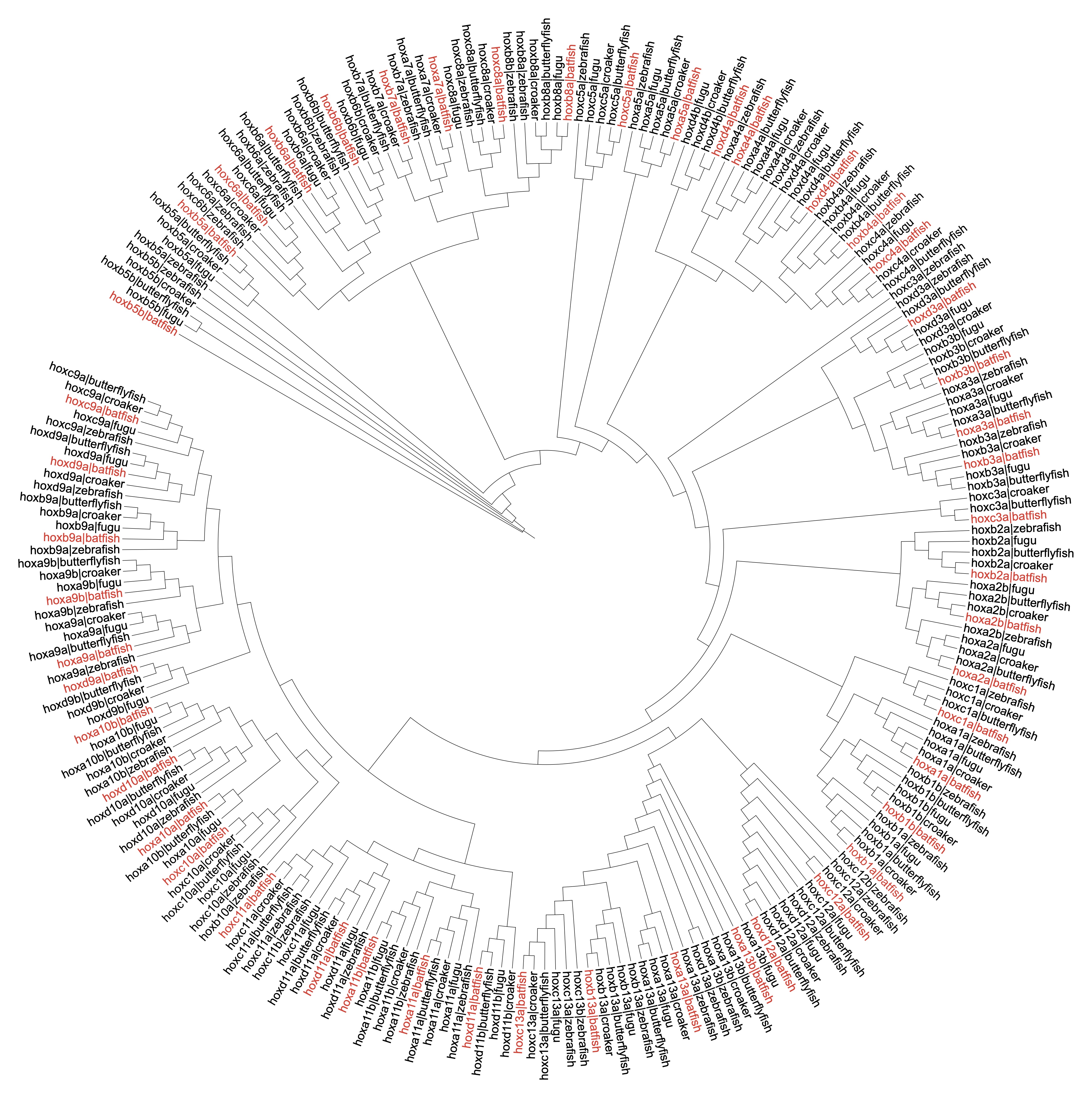

**Figure S5.** Phylogenetic tree of Hox proteins from orbicular batfish, copperband butterflyfish, large yellow croaker, Japanese pufferfish (fugu), and zebrafish. Hox proteins from orbicular batfish are highlighted in red.
